# Reduction of chemokine CXCL9 expression by omega-3 fatty acids via ADP-ribosylhydrolase ARH3 in MIN6 insulin-producing cells

**DOI:** 10.1101/2024.01.30.578079

**Authors:** Youngki You, Soumyadeep Sarkar, Cailin Deiter, Emily C. Elliott, Carrie D. Nicora, Raghavendra G. Mirmira, Lori Sussel, Ernesto S. Nakayasu

## Abstract

Type 1 diabetes (T1D) results from the autoimmune destruction of the insulin producing β cells of the pancreas. Omega-3 fatty acids protect β cells and reduce the incident of T1D. However, how omega-3 fatty acids act on β cells is not well understood. We have shown that omega-3 fatty acids reduce pro-inflammatory cytokine-mediated β-cell apoptosis by upregulating the expression of the ADP-ribosylhydrolase ARH3. Here, we further investigate the β-cell protection mechanism by ARH3 by performing siRNA of its gene *Adprhl2* in MIN6 insulin-producing cells followed by treatment with a cocktail of the pro-inflammatory cytokines IL-1β + IFN-γ + TNF-α, and proteomics analysis. ARH3 regulated proteins from several pathways related to the nucleus (splicing, RNA surveillance and nucleocytoplasmic transport), mitochondria (metabolic pathways) and endoplasmic reticulum (protein folding). ARH3 also regulated the levels of cytokine-signaling proteins related to the antigen processing and presentation, and chemokine-signaling pathway. We further studied the role of ARH in regulating the chemokine CXCL9. We confirmed that ARH3 reduces the cytokine-induced expression of CXCL9 by ELISA. We also found that CXCL9 expression is regulated by omega-3 fatty acids. In conclusion, we showed that omega-3 fatty acids regulate CXCL9 expression via ARH3, which might have a role in protecting β cells from immune attack and preventing T1D development.

## Introduction

Type 1 diabetes affects approximately 8.4 million people worldwide and it is cause by the autoimmune destruction of the insulin-producing β cells of the pancreas [1, 2]. This autoimmune response is characterized by infiltration of immune cells to the pancreatic islets, production of pro-inflammatory cytokines and chemokines, and circulating auto-antibodies against islet proteins [3]. The immune cells in conjunction with cytokines and chemokines induce apoptosis of β cells [3]. However, the signaling networks that control this autoimmune attack is not completely understood.

Protein post-translational modifications, such as phosphorylation, acetylation, and ubiquitination, are major regulators of signal transduction in cells [4]. ADP-ribosylation is a modification on which adenosine diphosphate (ADP)-ribose is transferred to proteins, RNAs and DNAs by ADP-ribosyltransferases (PARPs or ADRTs) using nicotinamide-adenosine dinucleotide (NAD) as a donor [5]. ADP-ribosylation can occur as single units (mono-ADP-ribosylation or MARylation) or chains (poly-ADP-ribosylation or PARylation) [5]. ADP-ribosylation regulates many processes of the immune response. For instance, several PARPs play central role in antiviral responses. ADP-ribosylation are key players of the stress granule formation, which sequester viral and cellular RNAs into cytosolic aggregates and halt their translation [6]. Moreover, ADP-ribosylation enhances the activity of transcription factors involved in inflammatory responses, such as STATs, NF-κB and NFATc3, upregulating the production of cytokines and chemokines [7-9].

Only few functions are known for ADP-ribosylation in T1D development. For instance, deletion of PARP1 protects mice from developing diabetes induced by streptozotocin [10-12]. Moreover, a variety of ADP-ribosyltransferases, including PARP3, -9, -10, -12, and - 14, are strongly regulated by pro-inflammatory cytokines in pancreatic islets [13]. We have found that the ADP-ribosylhydrolase ARH3, which cleaves ADP-ribosylation on serine residues [14], suppress apoptosis in MIN6 cells [13]. The mechanism includes degradation of SUZ12, a component of the histone methylation polycomb PCR2, induced by omega-3 fatty acids. SUZ12 degradation reduces histone methylation, increasing the expression of ARH3. The increase in ARH3 expression presumably protects cells by hydrolyzing ADP-ribosylation that are involved in pro-apoptotic signaling [13]. As mentioned, to this date very little is known about the signaling regulated by ARH3.

Here, we studied the role of ARH3 in regulating pro-inflammatory cytokine signaling. We performed proteomics of ARH3 knockdown MIN6 cells treated or not with the cocktail of pro-inflammatory cytokines IL-1β + IFN-γ + TNF-α. We also performed ELISA and qPCR to validate a protein regulated by ARH3 and to test the influence of omega-3 fatty acids on this regulation.

## Material and Methods

### MIN6 β cell line culture and treatment

MIN6 cells were cultured in DMEM containing 10% FBS, 50μM 2-mercaptoethanol and 1% penicillin-streptomycin and maintained at 37ºC in 5% CO_2_ atmosphere. For knockdown experiments, cells were transfected using Lipofectamine RNAiMAX (Invitrogen) with SMARTpool ON-TARGETplus non-targeting siRNA (Cat# D-001810-10-50, Dharmacon) as control or siRNA targeting the ARH3 gene *Adprhl2* (Cat# L-051819-01-0020, Dharmacon). To achieve robust knockdown of ARH3, cells were transfected a second time with NT or ARH3 siRNA 24 h after the first transfection. In experiments utilizing cytokines, cells were treated for 24 h with 100 ng/mL IFN-γ, 10 ng/mL TNF-α, and 5 ng/mL IL-1β.

### Proteomic analysis

For proteomics analysis, treated cells were rinsed with PBS before harvesting. Cells were dissolved in 50 mM NH_4_HCO_3_ containing 8 M urea and 10 mM dithiothreitol. After incubating for 1 h at 37 ºC with shaking at 800 rpm, 400 mM iodoacetamide was added to a final concentration of 40 mM, and the mixture incubated for another hour in the dark at room temperature. The reaction mixture was 8-folds diluted with 50 mM NH_4_HCO_3_, and 1 M CaCl_2_ was added to a final concentration of 1 mM. Proteins were digested for 3 h at 37 ºC using trypsin at 1:50 enzyme:protein ratio. Digested peptides were desalted by solid-phase extraction using C18 cartridges (Discovery, 50 mg, Supelco) and dried in a vacuum centrifuge.

Peptides were analyzed on a Waters NanoAquity UPLC system coupled with a Q-Exactive mass spectrometer (Thermo Scientific), as previous described in detail [13]. Data were processed with MaxQuant software (v1.6.14.0) [15] by matching against the mouse reference proteome database from Uniprot Knowledge Base (downloaded on August 31, 2020). Searching parameters included protein N-terminal acetylation and oxidation of methionine as variable modifications, and carbamidomethylation of cysteine residues as fixed modification. Mass shift tolerance was used as the default setting of the software. Only fully tryptic digested peptides were considered, allowing up to two missed cleaved sites per peptide. Identifications were filtered at 1% false-discovery rate in both peptide-spectrum match and protein group levels.

Quantitative information was extracted using the LFQ function of MaxQuant and analyzed in Perseus [16]. Proteins were considered differentially abundant with a p ≤ 0.05 using Student’s *t*-test. For functional-enrichment analysis, DAVID [17] analysis was used by querying the differentially abundant proteins against the entire mouse genome as background ontologies were considered enriched with a p ≤ 0.05 using Fisher’s exact test. The network of immune proteins was plotted with VANTED [18].

### ELISA

The levels of CXCL9 secreted to the media by MIN6 cells were quantified by Mouse DuoSet ELISA kit (R&D Systems). Polystyrene 96-well plates were coated with 100 μL of 200 ng/mL unconjugated goat anti-mouse CXCL9 antibody in PBS at room temperature overnight. Plates were washed 3x with 400 μL washing buffer and then blocked with 1% BSA for 1 h. After blocking, plates were washed 3x with 400 μL washing buffer. Cell culture supernatants were diluted 1:125 with reagent diluent and incubated for 2 h at room temperature, along with standards. Plates were washed 3x with 400 μL washing buffer and incubated with biotinylated goat anti-mouse CXCL9 antibody for 2 h at room temperature. Plates were washed 3x with 400 μL washing buffer and incubating with streptavidin-HRP incubation for 20 min at room temperature. Plates were washed 3x with 400 μL washing buffer, developed with tetramethylbenzidine and H_2_O_2_, and read at 570 nm in a plate reader (BioTek, USA).

### Quantitative real-time PCR analysis

Cells were harvested using Tri reagent (Zymo, Cat#R2050-1-200) and total mRNA was extracted using the RNA Clean & Concentrator™-5 (Zymo, Cat# R1014). Total RNA quantity was measured using nanodrop and qRT-PCR assays were performed using QuantiNova™ SYBR Green RT-PCR Kit (Qiagen, Cat.# ID: 208154) in StepOnePlus™ Real-Time PCR System by Applied biosystems. Expression levels of *Adprhl2* (R: CAAACTTCTGTACATCTTGGAC & F: AGAAACTCCTGAATCCCAAG) & *Cxcl9* (R: GTTTGATCTCCGTTCTTCAG & F: GAGGAACCCTAGTGATAAGG) were normalized to two housekeeping gene *Nono* (R: CATACTCATACTCAAAGGAGC & F: CTTCTTGCTGACTACATTTCC) & *Rpl13a* (R: CAGGTAAGCAAACTTTCTGG, F: CCTATGACAAGAAAAAGCGG).

## Results

### ARH3 regulated proteins in MIN6 cells

To investigate roles of ARH3 in cells, we performed siRNA of its gene *Adprhl2* (si*ARH3*) in MIN6 insulin-producing cells for 24 h, followed by a 24 h treatment with a cocktail of pro-inflammatory cytokines (IL-1β + IFN-γ + TNF-α) and submitted to proteomics analysis (**Figure 1A**). The analysis resulted in the identification and quantification of 4,749 proteins (**Supplemental Table 1**). The abundance of ARH3 was assessed by extracting the intensity-based absolute quantification (iBAQ) values with the MaxQuant software, which revealed a reduction in 82% in the ARH3 protein abundance (**Figure 1B**). We first analyzed the impact of si*ARH3* in cells without the cytokine treatment. The comparative analysis revealed that 159 and 161 proteins were up and downregulated with si*ARH3*, respectively (**Figure 1C**). An enrichment analysis of the subcellular localization revealed that the cytoplasm, nucleus, mitochondrion, and endoplasmic reticulum are the most common locations of the proteins regulated by ARH3 (**Figure 1D**). Further, a functional-enrichment analysis revealed an enrichment of proteins from a variety of pathways (**Figure 1E**). In agreement with the subcellular localization, the oxidative phosphorylation, Parkinson disease, prion disease, amyotrophic lateral sclerosis, Huntington disease, and thermogenesis pathways (**Figure 1E**) have the mitochondrion as one of their main components. In addition to oxidative phosphorylation and thermogenesis, the siARH3 regulated two other metabolic pathways, folate and pyrimidine metabolisms (**Figure 1E**). This indicates that ARH3 might have a role in regulating the mitochondrion and the cellular metabolism. Similar enrichment of pathways was observed in nuclear proteins, being the spliceosome, thermogenesis, and circadian entrainment pathways (**Figure 1E**) associated at least in part with this organelle. Protein processing in endoplasmic reticulum (ER), Parkinson disease, prion disease, amyotrophic lateral sclerosis, Huntington disease are pathways associated with endoplasmic reticulum (**Figure 1E**). Overall, the functional-enrichment analysis indicates that ARH3 regulates pathways that are associated with the mitochondrion, nucleus, and ER.

**Figure 1.**
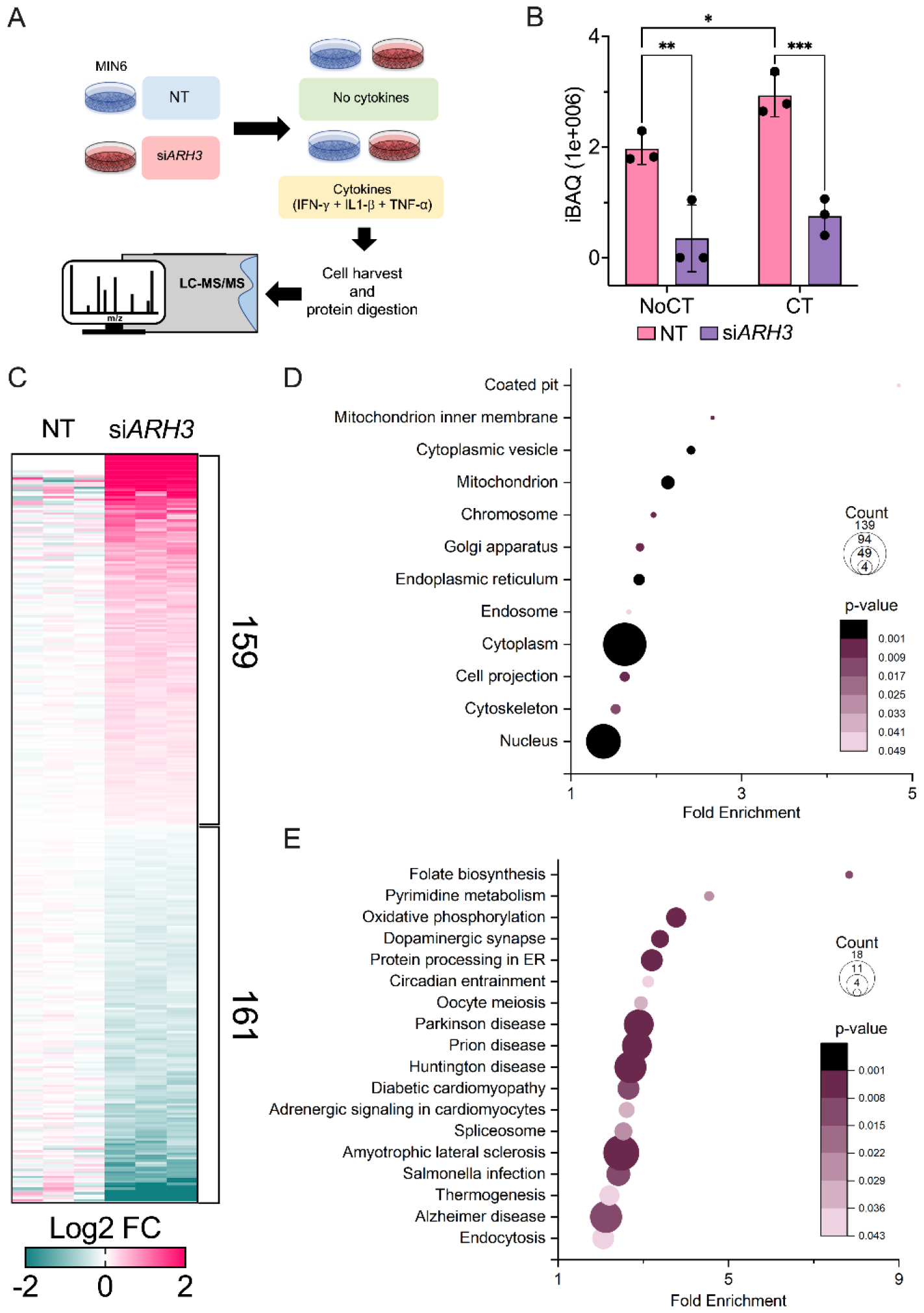
Proteomics analysis of ARH3 knockdown in MIN6 cells. (A) Experimental design. (B) Abundance of ARH3 protein detected in the proteomics analysis. Statistical significance was determined 2-way ANOVA with uncorrected Fisher’s least significant difference test: *p≤0.05, **p≤0.01, ***p≤0.001. (C) Heatmap of proteins differentially abundant in ARH3 knockdown vs. control MIN6 cells. This heatmap includes only samples from cells that were not treated with cytokines. (D) Enrichment of intracellular localization among the differentially abundant in ARH3 knockdown vs. control MIN6 cells using DAVID functional enrichment analysis. (E) Enrichment of pathways among the differentially abundant in ARH3 knockdown vs. control MIN6 cells using DAVID functional enrichment analysis. Abbreviations: CT – cocktail of pro-inflammatory cytokines IL-1β + IFN-γ + TNF-α, NoCT – control without the cytokine cocktail treatment, si*ARH3* – *Adprhl2* (ARH3 gene) siRNA.

### ARH3-regulated proteins in cytokine-treated cells

We study the impact of ARH3 in regulating proteins in cytokine-treated cells by comparing the parental and si*ARH3* MIN6 cells (**Figure 1A**). Confirming previous findings [13], ARH3 abundance increased 49% with the cytokine treatment (**Figure 1B**). Similar to the untreated cells, si*ARH3* reduced the abundance of ARH3 in 74% in cytokine-treated cells (**Figure 1B**). Our analysis showed that 158 and 359 proteins significantly were up and down regulated by si*ARH3*, respectively (**Figure 2A**). A functional-enrichment analysis showed a similar trend of the cytosol, nucleus, mitochondrion, and ER being the most abundant cellular localizations of the ARH3 regulated proteins in cytokine-treated cells (**Figure 2B**). However, the nucleus had a noticeable increase in the number of ARH3-regulated proteins, which increased from 44 (13.7% of the proteins assigned to specific cellular localizations) (**Figure 1C**) to 219 proteins (42% of the proteins assigned to specific cellular localizations) (**Figure 2B**). Mitochondrial and ER had similar proportion of proteins assigned to specific cellular localizations in both untreated (mitochondria – 13.7%, ER – 11.5%) and cytokine-treated cells (mitochondria – 13.4%, ER – 12.3%) (**Figures 1C** and **2B**). The increase in nuclear proteins regulated by ARH3 after cytokine-treated also reflected at the pathway-enrichment level. The spliceosome and thermogenesis pathways were similar regulated by ARH3 in untreated and cytokine-treated cells, although the number of regulated proteins increased after the cytokine treatment (**Figures 1D** and **2C**). In cells treated with cytokines, additional nuclear pathways were enriched among the ARH3-regulated proteins, including nucleocytoplasmic transport, mRNA surveillance, viral carcinogenesis pathways (**Figure 2C**). As histones are major ARH3 substrates [19], we looked at the levels of these proteins in the proteomics analysis. We found that the si*ARH3* reduced the levels of histone H1.5 in the cells treated with cytokines, and the levels of histone H1.2 and H2B in untreated cells (**Figure 3**). Together, these results show that in the presence of cytokines, ARH3 regulates a variety of proteins involved nuclear processes, such as splicing, nucleocytoplasmic transport and mRNA surveillance.

**Figure 2.**
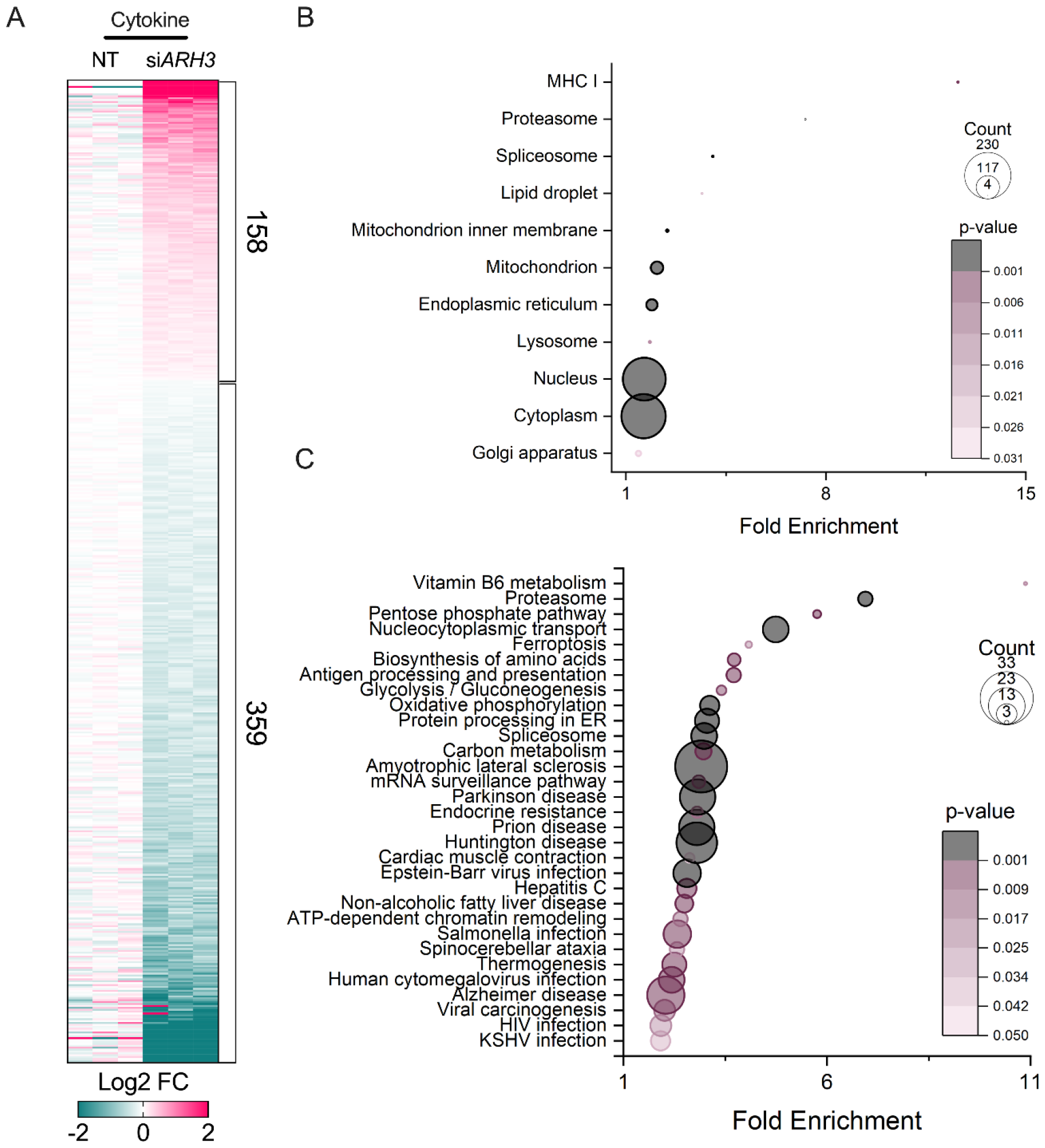
Proteomics analysis of ARH3-regulated proteins in MIN6 cells treated with the pro-inflammatory cytokines IL-1β + IFN-γ + TNF-α. (A) Heatmap of proteins differentially abundant in siARH3 vs. control MIN6 cells (NT – non targeted siRNA) treated with the cytokine cocktail. (B) Enrichment of subcellular localization among the differentially abundant proteins using DAVID functional enrichment analysis. (C) Enrichment of pathways among the differentially abundant proteins using DAVID functional enrichment analysis. Abbreviations: CT – cocktail of pro-inflammatory cytokines IL-1β + IFN-γ + TNF-α, NoCT – control without the cytokine cocktail treatment, si*ARH3* – *Adprhl2* (ARH3 gene) siRNA.

**Figure 3.**
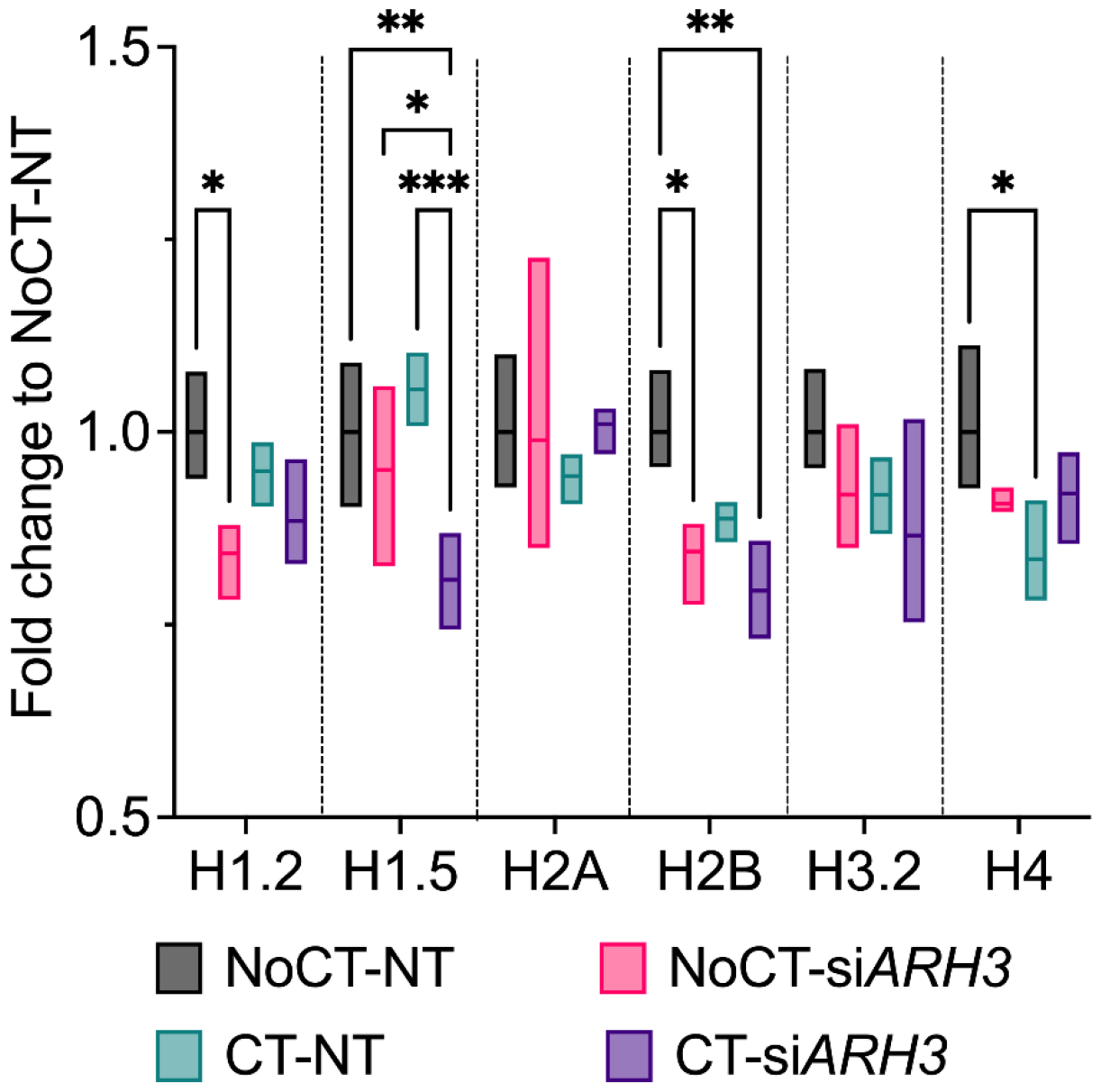
Quantification of histones in the proteomics analysis. Statistical significance was determined 2-way ANOVA with uncorrected Fisher’s least significant difference test: *p≤0.05, **p≤0.01, ***p≤0.001. Abbreviations: CT – cocktail of pro-inflammatory cytokines IL-1β + IFN-γ + TNF-α, NoCT – control without the cytokine cocktail treatment, si*ARH3* – *Adprhl2* (ARH3 gene) siRNA.

### Regulation of immune proteins by both cytokine and ARH3 signaling

We focus next on the proteins that were significantly regulated by the cytokine cocktail and were further regulated by si*ARH3*. A total of 128 proteins were significantly regulated by both the cytokine treatment and siARH3 (**Figure 4A**). The functional enrichment analysis revealed that these proteins were enriched in a variety of immune-related pathways, such as antigen processing and presentation (including proteasome), chemokine signaling pathway, and infections (Epstein-Barr virus, human cytomegalovirus, human immunodeficiency virus, toxoplasmosis, Kaposi sarcoma-associated herpesvirus and viral carcinogenesis) (**Figure 4B**). To further investigate how the regulation and connection of these 128 regulated proteins, we extracted the interaction networks from the String database. We found 18 inter-connected proteins from the chemokine signaling pathway and immune system pathway (**Figure 4C**). The levels of the cytokine-induced transcription factors Stat1 and Stat3 had a small but significant reduction with si*RNA3* (**Figure 4C**). Similar trend was observed for the proteins involved in antigen processing and presentation, H2-K1, B2m, Tap2, Psmb8, Psmc2, Psmb10 and Hsp90ab1 (**Figure 4C**), possibly in response to Stat1 and Stat3 regulation. Conversely, the levels of the autophagy receptor Sqstm1, nitric oxide synthase Nos2, and chemokine Cxcl9 were upregulated in siARH3 (**Figure 4C**). These results show that ARH3 regulates cytokine-mediated signaling, including the levels of its main transcription factors Stat1 and Stat3, along with other immune proteins.

**Figure 4.**
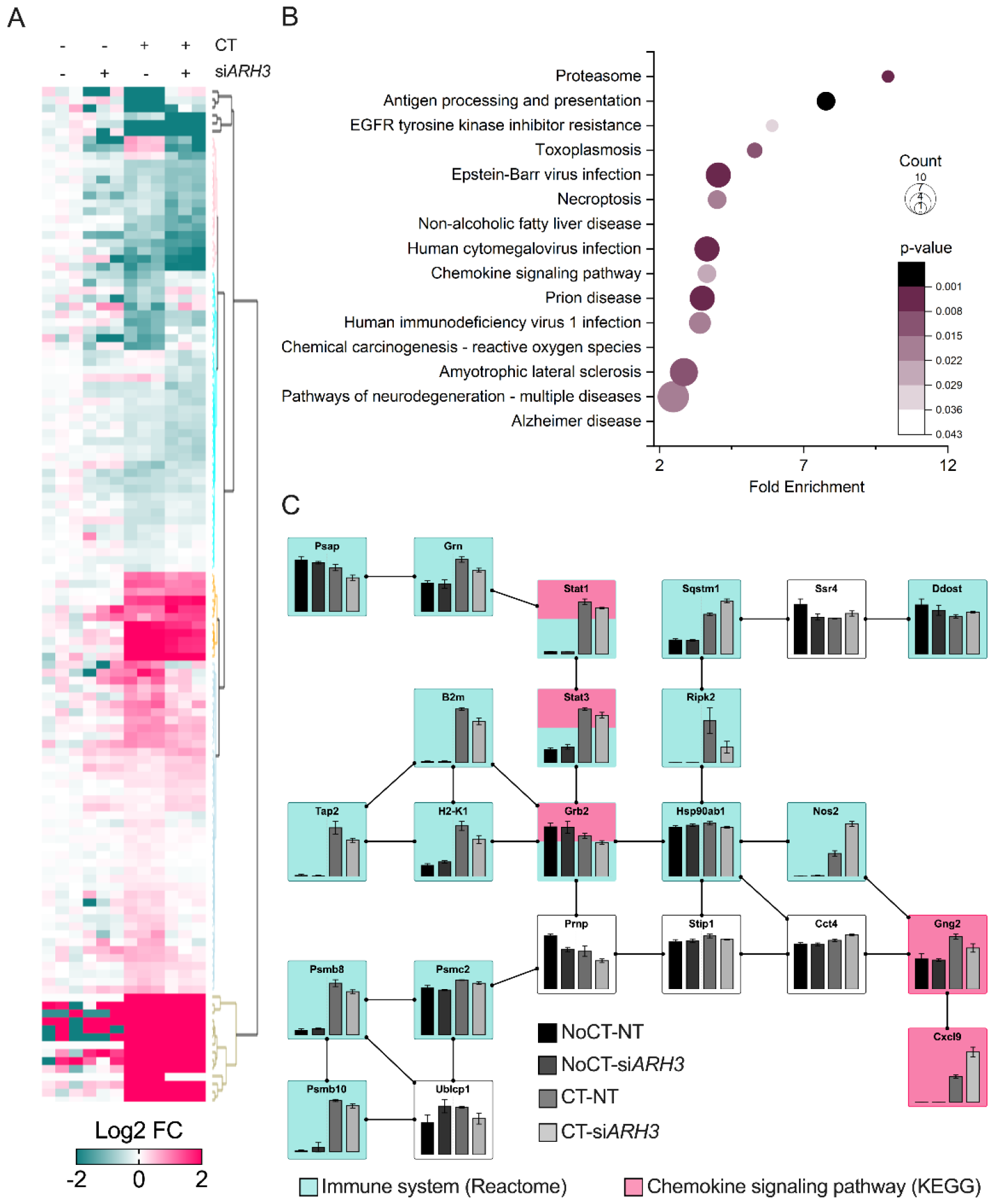
Proteins regulated by both cytokine and ARH3-mediated signaling. (A) Heatmap of proteins differentially abundant in both comparisons, cytokine-treated vs. untreated non-targeted siRNA MIN6 cells and siARH3 vs. non-targeted siRNA in cells treated with cytokines. (B) Enrichment of pathways among the differentially abundant proteins using DAVID functional enrichment analysis. (C) The network of regulated immune proteins extracted from KEGG and Reactome databases and plotted along their relative abundance (normalized by the highest) using VANTED. Abbreviations: CT – cocktail of pro-inflammatory cytokines IL-1β + IFN-γ + TNF-α, NoCT – control without the cytokine cocktail treatment, si*ARH3* – *Adprhl2* (ARH3 gene) siRNA.

We further studied possible mechanisms of the chemokine CXCL9 expression regulation by ARH3. We validate the regulation of CXCL9 by ELISA of the supernatant from cell cultures, which confirmed that this protein is induced by the pro-inflammatory cytokine cocktail and that the si*ARH3* further increased its production (**Figure 5A**). As we previously found that ARH3 expression is regulated by omega-3 fatty acids [13], we asked the question if this regulation would influence the expression of CXCL9. Therefore, we pre-treated MIN6 cells with the omega-3 fatty acids docosahexaenoic acid and eicosapentaenoic acid in combination with the cytokine cocktail for 8 h. The levels of CXCL9 transcript were measured by qPCR. The expression of CXCL9 was induced by the cytokine cocktail, which was reduced in 38.6% and 44.4% by eicosapentaenoic acid and docosahexaenoic acid, respectively (**Figure 5B**). These results show that ARH3 downregulates the expression/production of CXCL9, which is likely to be via ARH3 expression regulation by omega-3 fatty acids.

**Figure 5.**
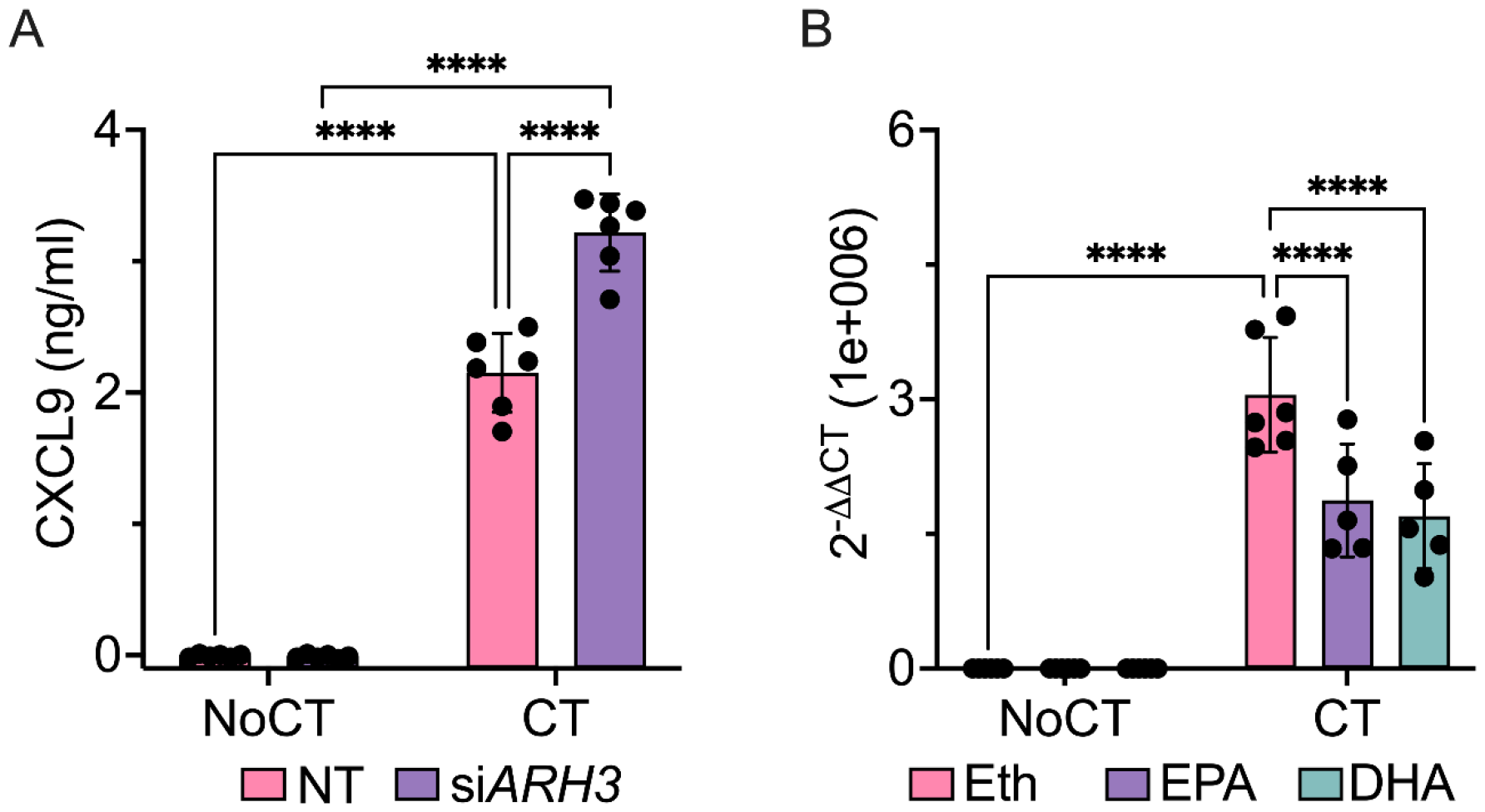
Regulation of CXCL9 by ARH3 and omega-3 fatty acids. (A) Quantification of CXCL9 secreted into the culture media by ARH3 knockdown (siARH3) MIN6 cells treated or not with the pro-inflammatory cytokines IL-1β + IFN-γ + TNF-α. (B) Quantification of *Cxcl9* gene expression in MIN6 cells treated or not with the pro-inflammatory cytokines IL-1β + IFN-γ + TNF-α in combination with omega-3 fatty acids (DHA and EPA). Statistical significance was determined 2-way ANOVA with uncorrected Fisher’s least significant difference test: *p≤0.05, **p≤0.01, ***p≤0.001. Abbreviations: CT – cocktail of pro-inflammatory cytokines IL-1β + IFN-γ + TNF-α, DHA – docosahexaenoic acid, EPA -eicosapentaenoic acid EPA, Eth – ethanol (solvent vehicle), NoCT – control without the cytokine cocktail treatment, si*ARH3* – *Adprhl2* (ARH3 gene) siRNA.

## Discussion

Here we investigate the roles of ARH3 in MIN6 cells by proteomics analysis. Our data show that ARH3 regulates pathways associated with mitochondrion, nucleus, and ER. This agrees with previous observations that ARH3 hydrolyses ADP-ribosylation from both mitochondrial and nuclear proteins [20, 21]. The regulation of mitochondrial proteins by ARH3 was accompanied with enrichment in metabolic pathways in both untreated and cytokine treated MIN6 cells. However, the regulation of specific metabolic pathways by ARH3 has not been reported yet and needs further studies. The regulation of mitochondrial proteins by ARH3 may play role in apoptosis, since we have previous described that this protein reduces apoptosis in MIN6 cells and mitochondrion is a main organelle involved in this process. In addition, ADP-ribosylation has also been described to play an important role in apoptosis [22]. In the ADP-ribosylation database ADPriboDB 2.0 [23], several apoptotic signaling proteins have been listed as being ADP-ribosylated, including FADD, Casp8, BID, BCL2, BCL2L1, Bax, Cycs, Diablo, BIRC2 and BIRC6. More recently, the apoptotic proteins API5, ACIN1, CCAR1, PARW, AATF and CIAPIN1, were found to be ARH3 substrates [19]. However, the strongest evidence so far is that ARH3 regulates apoptotic signaling by preventing the release of apoptosis-inducing factor AIF from mitochondria [21]. Therefore, more investigation is required to have a better understanding of the ARH3 functions in the mitochondrion. Our data also show that ARH3 regulates variety of proteins from pathways associated with the ER. To the best of our knowledge, no functions have been associated with ARH3 in this organelle.

Our proteomics data also show that nuclear ARH3-regulated proteins were enriched in RNA-related pathways, such as splicing, nucleocytoplasmic transport, mRNA surveillance. ARH3 can hydrolyze ADP-ribosylation from RNA chains [14, 24], but its function is still poorly understood. However, ADP-ribosylation itself is well described to regulate RNA biology, including all forementioned pathways [25]. Therefore, it is possible that ARH3 is a major regulator ADP-ribosylation-mediated RNA regulation in cells. Our data also show that ARH3 regulates proteins involved in gene expression, such as histones and the transcription factors Stat1 and Stat3. The reduction of Stat1 and Stat3 in si*ARH3* cells might be related to the downregulation of antigen processing and presentation proteins. Stat1 and Stat3 are transcription factors of antigen processing and presentation proteins, and they were regulated in similar levels. The reduction in histone levels in si*ARH3* cells might be a measurement artifact of the reduced detectability of the ADP-ribosylated peptide in the mass spectrometer. ADP-ribosylation in histones is well described and ARH3 is a major regulatory enzyme of this process [19]. ADP-ribosylation of histones are associated with chromatin opening and increase in gene expression, including cytokine and chemokine genes [22]. Therefore, ARH3-regulation of histone ADP-ribosylation could be associated with CXCL9 expression. The inflammatory transcription factor NF-κB activity is enhanced by PARP1, not by ADP-ribosylation of itself [8]. It is possible that NF-κB activity is increased by ADP-ribosylation of other proteins, such as histones. Another possibility is that ARH3 is removing ADP-ribosylation from Stat1 and repressing its activity [26], as this transcription factor is a major activator of CXCL9 gene expression in β cells [7, 9].

We also showed that reduction of cytokine-induced CXCL9 expression by ARH3 involves omega-3 fatty acids. We previously showed that omega-3 fatty acids improves the expression of ARH3 by reducing the level of Suz12, a component of the histone methylation polycomb PCR2 [13]. Omega-3s are anti-inflammatory fatty acids that regulate the expression of pro-inflammatory cytokines and chemokines by engaging the G protein coupled receptor GPR120, inhibiting the activity of the pro-inflammatory transcription factor NF-κB, and activating the anti-inflammatory transcription factor peroxisome proliferator activated receptor γ [27]. The signal transduction of the GPR120 involves the sequestration of the NF-κB activator Tab1, but this scenario is not completely understood.

In conclusion, in this paper we found that ARH3 regulates a variety of proteins and pathways, in special, the ones related to the cell mitochondria and nuclei. We show that ARH3 reduces the expression of the chemokine CXCL9 in response to cytokine signaling, which is mediated by omega-3 fatty acids.

## FUNDING

This work was supported by the National Institute of Diabetes and Digestive and Kidney Diseases grants U01 DK127505 (to L.S. and E.S.N.), U01 DK127786 (to R.G.M.), R01 DK060581 (to R.G.M.). E.S.N. was also supported by the Catalyst Award from the Human Islet Research Network.

## ACKNOWLEDGMENTS

The authors thank Drs. Charles Ansong and Charanya Muralidharan for their insightful inputs. Part of the work was performed in the Environmental Molecular Sciences Laboratory, a U.S. Department of Energy (DOE) national scientific user facility at Pacific Northwest National Laboratory (PNNL) in Richland, WA. Battelle operates PNNL for the DOE under contract DE-AC05-76RLO01830.

## CONFLICT OF INTEREST STATEMENT

The authors have declared no conflicts of interest.

## SUPPORTING INFORMATION

**Supplemental Table 1**. Proteomics analysis ARH3 knockdown in MIN6 cells treated with the cocktail of pro-inflammatory cytokines IL-1β + IFN-γ + TNF-α.

## Notes

### Competing Interest Statement

The authors have declared no competing interest.

## References

[1] Gregory, G. A., Robinson, T. I. G., Linklater, S. E., Wang, F., et al., Global incidence, prevalence, and mortality of type 1 diabetes in 2021 with projection to 2040: a modelling study. Lancet Diabetes Endocrinol 2022, 10, 741–760.

[2] DiMeglio, L. A., Evans-Molina, C., Oram, R. A., Type 1 diabetes. Lancet 2018, 391, 2449–2462.

[3] Eizirik, D. L., Pasquali, L., Cnop, M., Pancreatic beta-cells in type 1 and type 2 diabetes mellitus: different pathways to failure. Nat Rev Endocrinol 2020, 16, 349–362.

[4] Liu, J., Qian, C., Cao, X., Post-Translational Modification Control of Innate Immunity. Immunity 2016, 45, 15–30.

[5] Suskiewicz, M. J., Prokhorova, E., Rack, J. G. M., Ahel, I., ADP-ribosylation from molecular mechanisms to therapeutic implications. Cell 2023, 186, 4475–4495.

[6] Jayabalan, A. K., Adivarahan, S., Koppula, A., Abraham, R., et al., Stress granule formation, disassembly, and composition are regulated by alphavirus ADP-ribosylhydrolase activity. Proc Natl Acad Sci U S A 2021, 118.

[7] Nie, Y., Nirujogi, T. S., Ranjan, R., Reader, B. F., et al., PolyADP-Ribosylation of NFATc3 and NF-kappaB Transcription Factors Modulate Macrophage Inflammatory Gene Expression in LPS-Induced Acute Lung Injury. J Innate Immun 2021, 13, 83–93.

[8] Hassa, P. O., Hottiger, M. O., The functional role of poly(ADP-ribose)polymerase 1 as novel coactivator of NF-kappaB in inflammatory disorders. Cell Mol Life Sci 2002, 59, 1534–1553.

[9] Iwata, H., Goettsch, C., Sharma, A., Ricchiuto, P., et al., PARP9 and PARP14 cross-regulate macrophage activation via STAT1 ADP-ribosylation. Nat Commun 2016, 7, 12849.

[10] Andreone, T., Meares, G. P., Hughes, K. J., Hansen, P. A., Corbett, J. A., Cytokine-mediated beta-cell damage in PARP-1-deficient islets. Am J Physiol Endocrinol Metab 2012, 303, E172–179.

[11] Burkart, V., Wang, Z. Q., Radons, J., Heller, B., et al., Mice lacking the poly(ADP-ribose) polymerase gene are resistant to pancreatic beta-cell destruction and diabetes development induced by streptozocin. Nat Med 1999, 5, 314–319.

[12] Masutani, M., Suzuki, H., Kamada, N., Watanabe, M., et al., Poly(ADP-ribose) polymerase gene disruption conferred mice resistant to streptozotocin-induced diabetes. Proc Natl Acad Sci U S A 1999, 96, 2301–2304.

[13] Sarkar, S., Deiter, C., Kyle, J. E., Guney, M. A., et al., Regulation of β-cell death by ADP-ribosylhydrolase ARH3 via lipid signaling in insulitis. Cell Commun Signal, In press.

[14] Abplanalp, J., Leutert, M., Frugier, E., Nowak, K., et al., Proteomic analyses identify ARH3 as a serine mono-ADP-ribosylhydrolase. Nat Commun 2017, 8, 2055.

[15] Cox, J., Mann, M., MaxQuant enables high peptide identification rates, individualized p.p.b.-range mass accuracies and proteome-wide protein quantification. Nat Biotechnol 2008, 26, 1367–1372.

[16] Tyanova, S., Temu, T., Sinitcyn, P., Carlson, A., et al., The Perseus computational platform for comprehensive analysis of (prote)omics data. Nat Methods 2016, 13, 731–740.

[17] Huang da, W., Sherman, B. T., Lempicki, R. A., Systematic and integrative analysis of large gene lists using DAVID bioinformatics resources. Nat Protoc 2009, 4, 44–57.

[18] Junker, B. H., Klukas, C., Schreiber, F., VANTED: a system for advanced data analysis and visualization in the context of biological networks. BMC Bioinformatics 2006, 7, 109.

[19] Hendriks, I. A., Buch-Larsen, S. C., Prokhorova, E., Elsborg, J. D., et al., The regulatory landscape of the human HPF1- and ARH3-dependent ADP-ribosylome. Nat Commun 2021, 12, 5893.

[20] Demichev, V., Messner, C. B., Vernardis, S. I., Lilley, K. S., Ralser, M., DIA-NN: neural networks and interference correction enable deep proteome coverage in high throughput. Nat Methods 2020, 17, 41–44.

[21] Mashimo, M., Kato, J., Moss, J., ADP-ribosyl-acceptor hydrolase 3 regulates poly (ADP-ribose) degradation and cell death during oxidative stress. Proc Natl Acad Sci U S A 2013, 110, 18964–18969.

[22] Kunze, F. A., Hottiger, M. O., Regulating Immunity via ADP-Ribosylation: Therapeutic Implications and Beyond. Trends Immunol 2019, 40, 159–173.

[23] Ayyappan, V., Wat, R., Barber, C., Vivelo, C. A., et al., ADPriboDB 2.0: an updated database of ADP-ribosylated proteins. Nucleic Acids Res 2021, 49, D261–D265.

[24] Weixler, L., Feijs, K. L. H., Zaja, R., ADP-ribosylation of RNA in mammalian cells is mediated by TRPT1 and multiple PARPs. Nucleic Acids Res 2022, 50, 9426–9441.

[25] Kim, D. S., Challa, S., Jones, A., Kraus, W. L., PARPs and ADP-ribosylation in RNA biology: from RNA expression and processing to protein translation and proteostasis. Genes Dev 2020, 34, 302–320.

[26] Luo, X., Ryu, K. W., Kim, D. S., Nandu, T., et al., PARP-1 Controls the Adipogenic Transcriptional Program by PARylating C/EBPbeta and Modulating Its Transcriptional Activity. Mol Cell 2017, 65, 260–271.

[27] Diaz Ludovico, I., Sarkar, S., Elliott, E., Virtanen, S. M., et al., Fatty acid-mediated signaling as a target for developing type 1 diabetes therapies. Expert Opin Ther Targets 2023, 27, 793–806.

